# Psilocybin induces spatially constrained alterations in thalamic functional organizaton and connectivity

**DOI:** 10.1101/2022.02.28.482395

**Authors:** Andrew Gaddis, Daniel E. Lidstone, Mary Beth Nebel, Roland Griffiths, Stewart H. Mostofsky, Amanda Mejia, Frederick Barrett

**Author notes:** These authors contributed equally to this manuscript (Andrew Gaddis, Daniel E. Lidstone). **Correspondence:** Andrew Gaddis, Frederick Barrett.

## Abstract

**Background:** Serotonin 2A receptor (5-HT_2AR_) agonist psychedelics including psilocybin and LSD (“classic” psychedelics) evoke acute alterations in perception and cognition. Altered thalamocortical connectivity has been proposed to underlie these effects, which is supported by some functional MRI (fMRI) studies. Likely due to sample size limitations, these studies have treated the thalamus as a unitary structure, despite known differential 5-HT_2AR_ expression and functional specificity of different intrathalamic nuclei. Independent Component Analysis (ICA) has been employed to generate functional subdivisions of the thalamus from resting state fMRI (rsfMRI) data. This report utilizes a novel data-sparing ICA approach in order to examine psilocybin-induced changes in intrathalamic functional organization and thalamocortical connectivity.

**Methods:** Baseline rsfMRI data (n=38) was utilized to generate a template, which was then applied in a novel ICA-based analysis of the acute effects of psilocybin on intra- and extra-thalamic functional organization and connectivity in a smaller sample (n=18). Correlations with subjective reports of drug effect and comparisons with a previously reported analytic approach (treating the thalamus as a single functional unit) were conducted.

**Results:** Several intrathalamic components showed significant psilocybin-induced alterations in intrathalamic spatial organization, largely localized to the mediodorsal and pulvinar nuclei, and correlated with reported subjective effects. These same components demonstrated alterations in thalamocortical connectivity, largely with visual and default mode networks. Analysis in which the thalamus is treated as a singular unitary structure showed an overall numerical increase in thalamocortical connectivity, consistent with previous literature using this approach, but this increase did not reach statistical significance.

**Conclusions:** Utilization of a novel analytic approach demonstrated changes in intra- and extra-thalamic functional organization and connectivity of intrathalamic nuclei and cortical networks known to express the 5-HT_2AR_. Given that these changes were not observed using whole-thalamus analyses, it seems that psilocybin may cause widespread but modest increases in thalamocortical connectivity that are offset by strong focal decreases in functionally relevant intrathalamic nuclei.

## 1. INTRODUCTION

Psychedelic compounds can induce perceptual (Barrett et al., 2018b; Kometer and Vollenweider, 2018) and cognitive effects (Barrett et al., 2018a; Pokorny et al., 2020) that may be deeply impactful to the individuals experiencing them (Griffiths et al., 2008), and these compounds have potential utility in the treatment of psychiatric and neurologic disorders (Johnson et al., 2019). The perceptual and cognitive effects of “classic” psychedelics (e.g. LSD, psilocybin, mescaline) are thought to be mediated primarily through activation of the serotonin 2A receptor (5-HT_2AR_) (López-Giménez and González-Maeso, 2018; Nichols, 2016). This receptor is expressed widely across both cortical and subcortical regions (Beliveau et al., 2017; Hawrylycz et al., 2012); however the majority of research into the impact of psychedelic compounds on the brain have focused on regions expressing 5-HT_2AR_ in the cortex.

The thalamus is among the subcortical structures that demonstrate significant expression of 5-HT_2AR_ (Wai et al., 2011), and the perceptual and cognitive effects of the “classic” psychedelics have been hypothesized to be in part due to changes in cortico–striato–thalamo–cortical circuits that have also been implicated in the pathophysiology of psychosis (Geyer and Vollenweider, 2008; Vollenweider and Geyer, 2001; Vollenweider and Preller, 2020). Four converging lines of evidence support this hypothesis: 1) the 5-HT_2AR_ is expressed in the thalamus and other points within cortico–striato–thalamo-cortical circuits (Beliveau et al., 2017; Doss et al., 2021; Hawrylycz et al., 2012), 2) psychedelics have historically been framed as “psychotomimetic”, or sharing some elements of psychosis (Vollenweider et al., 1998); 3) there is a growing literature demonstrating that thalamocortical connectivity plays a key role in psychosis(Bergé et al., 2020; Hua et al., 2019) and, 4) there is growing evidence that thalamocortical connectivity plays a key role in both psychedelic and other altered perceptual and cognitive states with potential clinical relevance (Vollenweider & Preller, 2020).

The thalamus has long been recognized as a crucial “relay center” through which sensory information reaches the cortex. Consistent with this, altered gating of sensory information from primary sensory organs to the cortex by the thalamus (Liang et al., 2020; Sherman and Guillery, 2002) has been proposed as a potential mechanism behind altered perceptual states occurring during psychosis (Bergé et al., 2020; Geyer and Vollenweider, 2008; Vollenweider et al., 1998; Vollenweider and Geyer, 2001; Yao et al., 2020) and under anesthesia (White and Alkire, 2003). In recent decades it is increasingly recognized that the thalamus is also involved in the mediation and regulation of cortical signaling beyond sensory input (Sherman, 2007), with implications for involvement in cognitive processes such as attention and awareness, among others (De Bourbon-Teles et al., 2014). Changes in thalamocortical connectivity might also have clinical relevance in cognitive processes and states less directly related to sensory perception, such as meditation (Chen et al., 2021) and hypnosis (Huber et al., 2014). These studies collectively suggest a framework for understanding how changes in cortico–striato–thalamo–cortical connectivity might mediate the effects of psychedelic compounds, and how these changes could potentially play a role in the clinical utility of psychedelics.

To date, two fMRI studies in humans have reported differences in thalamic function during the acute effect of the classic psychedelic psilocybin. One found an overall decrease in cerebral blood flow to the thalamus (Carhart-Harris et al., 2012), and the other found an overall increased functional connectivity between the thalamus and a “task-positive” cortical network (Carhart-Harris et al., 2013). Differences in thalamic function under the acute influence of LSD, another classic psychedelic, have been more thoroughly characterized using fMRI in humans, with recent studies suggesting potential decreases as well as increases in thalamocortical connectivity. Two early studies reported increased global thalamic connectivity after administration of LSD (Müller et al., 2017; Tagliazucchi et al., 2016). A subsequent study suggested that visual, auditory, sensorimotor, default mode, and frontoparietal networks show both increases and decreases in connectivity with different thalamic regions during the acute effects of LSD (Müller et al., 2018). Another study employed global signal regression and reported a mix of increased and decreased thalamocortical connectivity after administration of LSD, with decreased thalamo-cortical connectivity in associative regions and increased thalamo-cortical connectivity in somatosensory regions (Preller et al., 2018). An even more recent study utilizing dynamic causal modeling demonstrated increased thalamic efferent connectivity to the posterior cingulate cortex and ventral striatum after acute LSD administration, with decreases in reciprocal afferent connections to the thalamus from these regions (Preller et al., 2019). Although consensus among the currently available literature seems to suggest a coarse large-scale increase in thalamic activation and thalamocortical connectivity under the influence of the classic psychedelics, the effect appears to be nuanced, with some reports utilizing alternative analytic approaches to demonstrate concurring decreases in connectivity.

Thalamic function is diverse, and the thalamus is widely understood to consist of numerous structurally and functionally distinct nuclei (Iglehart et al., 2020). Some thalamic nuclei demonstrate significant reciprocal connectivity with higher-order cortical regions, with each nucleus having dissociable connectivity and function (Halassa and Sherman, 2019). Animal models have demonstrated that there is reciprocal connectivity between cortical areas and thalamic nuclei that are known to express the 5-HT_2AR_ (Barre et al., 2016), suggesting that 5-HT_2AR_ signaling may alter the function of these circuits. Despite this, the thalamus is commonly treated as a unitary structure in functional neuroimaging studies with psychedelics. Practical limitations including small sample size and limited image resolution likely contribute to the persistence of whole-thalamus masking in analysis of thalamocortical connectivity via functional imaging, with sample size being a common limiting factor in psychedelic neuroimaging research.

In this study, we employ a hierarchical Template Independent Component Analysis (tICA) model of resting-state fMRI (rsfMRI) to estimate functional subdivisions of the thalamus, and then examine psilocybin-induced changes in thalamic functional organization and thalamocortical connectivity. ICA is a method that separates multivariate signals into distinct subcomponents, and has previously been applied to the subcortex and the thalamus, resulting in reliable identification of functionally distinct subregions in these areas (Tian et al., 2020). “Template” ICA is a recently developed data-efficient framework for ICA that leverages empirical population priors to optimize the estimation of functionally relevant regions in individual participants with standard-length fMRI scans (Mejia et al., 2020). Given the known anatomical and functional segregation of thalamic nuclei, this ICA-based approach provides an optimal analytic framework for further examination of the previously identified alterations in thalamocortical connectivity that occur during the acute effects of classic psychedelics, but on a level of resolution that may more faithfully reflect the differential function of independent thalamic nuclei.

## 2. METHODS

### 2.1 Participants

Participants (N=40) were enrolled in a multi-phase study of the acute and enduring subjective and neural effects of psilocybin in individuals with a long-term meditation practice (24 male/16 female: mean age 56 years old). All participants provided written informed consent, were medically and psychologically healthy as assessed by medical history and physical examination as well as the Structured Clinical Interview for DSM-IV (SCID-IV) and had a long-term meditation practice, defined as currently meditating for several years (e.g., 5 years) on a regular basis (e.g., at least 3 times per week; daily practice preferred), with experience using a range of different meditation techniques (e.g., concentration; compassion/loving-kindness; open awareness), and having attended at least 1 meditation retreat of 5 days or longer, or several weekend retreats. Participants enrolled in the study had average lifetime meditation hours of 4883 hours (range 1000-17000; STD = 4892).

Participants were excluded if they presented with clinically relevant cardiac abnormalities, had a first or second degree relative with a history of bipolar disorder, psychosis, or a related disorder, met criteria for substance use disorder, had a standard MRI contraindication (e.g. contraindicated medical devices or metal in the body), or had used a psychedelic drug after the establishment of their meditation practice. Half (50%) of the participants had previous experience with a hallucinogen before participating in the first phase of the experiment, but none had exposure to a hallucinogen within 5 years of study enrollment in the first phase of the experiment. Of those individuals who had previous hallucinogen exposure, mean previous hallucinogen use was 2.65 separate occasions (range = 1 to 20). All participants were confirmed negative by urinalysis for pregnancy (females) and for recent illicit drug use before enrollment and on the morning of drug administration and scanning procedures.

All procedures were approved by the Johns Hopkins Medicine Institutional Review Board and were carried out in accordance with the Declaration of Helsinki. This study was conducted as part of a clinical trial that was registered at ClinicalTrials.gov (NCT02145091). No serious adverse events were encountered, there were no cases of prolonged effects of drug administration, and no pharmacological, medical, or psychological interventions were necessary in response to study procedures.

### 2.2 Experimental Design

#### 2.2.1 Phase 1 Study Procedures

Forty healthy volunteers were enrolled in the first phase of this study, which consisted of screening and a baseline MRI procedure (prior to any drug administration), followed by administration of 25 mg/70 kg oral psilocybin two to four months later. Our template generation analysis is restricted to the restingstate functional MRI data collected during the baseline visit. These data have not been reported elsewhere. All participants completed 8 hours of meetings with study staff in *Phase 1* to build rapport and prepare for psilocybin administration (Johnson et al., 2008) in advance of the experimental drug administration session. By the end of *Phase 1*, all participants were familiar both with the MRI environment used in the current study and with the subjective effects of a high dose of psilocybin (25 mg/70 kg).

#### 2.2.2 Phase 2 Study Procedures

Twenty healthy volunteers from *Phase 1* of the study were enrolled in *Phase 2*, which consisted of a single-blind, within-subjects, placebo-controlled study of the acute effects of 10 mg/70 kg oral psilocybin on brain function. *Phase 2* was conducted approximately two months after the administration of 25 mg/70 kg oral psilocybin in *Phase 1* for each volunteer. Drug administration procedures are described below (see Section 2.2.3). Task-free functional MRI data from *Phase 2* were collected during both placebo and psilocybin conditions for each volunteer. Two volunteers were excluded based on unacceptable registration, and thus a total of 38 volunteers from Phase 1 and 18 volunteers from Phase 2 were included in the analysis (13M/5F; mean age = 54.4, SD = 13.2). Data from *Phase 2* have been analyzed and published elsewhere (Barrett et al., 2020).

In *Phase 2*, participants completed 4 hours of preparation before their drug administration sessions to reacclimate themselves to expected psilocybin effects and to prepare for experiencing psilocybin effects within the MRI. During sessions in *Phase 2*, participants were accompanied continuously, including in the MRI room, by at least one staff member, and were closely monitored during acute effects of placebo and psilocybin. After resolution of acute psilocybin effects and before leaving the research unit, all participants completed a debriefing with study staff.

#### 2.2.3 Psilocybin Scanning Procedures

To minimize expectancy effects, participants were informed both verbally and in the consent form that on each occasion, they may receive placebo, an extremely low dose of psilocybin, or a moderately low but still psychoactive dose of psilocybin (relative to the high dose of 25 mg/70 kg psilocybin they had received after their baseline MRI scan in Phase 1). Unknown to the participant, the first drug administration was always placebo (administered at 8am), and the second was always 10 mg oral psilocybin (administered at 12 noon). Psilocybin administration occurred in an aesthetic living-room environment in the Center for Psychedelic and Consciousness Research (CPCR) at the Johns Hopkins Bayview Medical Center, in Baltimore, MD. After capsule administration, participants reclined on a couch with eye shades and headphones and listened to music. Participants were transported by hired car and accompanied by two session monitors from the CPCR to the MRI facilities 60 min after capsule administration. Participants then completed scanning procedures beginning 90 min after administration of placebo or 10 mg/70 kg psilocybin. Task-free scans began approximately 100 min after capsule administration (±11 min). Scans were acquired on 3T Philips Achieva MRI scanners (Philips Medical Systems, Best, The Netherlands) at the F.M. Kirby Research Center for Functional Brain Imaging at the Kennedy Krieger Institute in Baltimore, MD. Each resting-state scan was acquired with eyes closed using echo planar imaging (axial acquisition, 3 mm isotropic voxel size, 1 mm slice gap, 37 ascending slices, 210 volumes retained for analysis, TR of 2.2s, TE of 30 ms, sense acceleration factor of 2, and four initial TRs dis-carded for magnetization equilibrium; total analyzed scan time was 7 min and 42s). The timing of scanning procedures coincided with peak subjective effects (Griffiths et al., 2006, 2011; Carbonaro et al., 2018) and expected peak plasma concentration of this dose of psilocybin (Brown et al., 2017). Participants were instructed to not meditate during task-free scans, and had experience completing resting-state scans following this instruction on at least one previous occasion. All participants confirmed after the MRI that they had not been meditating during the task-free scan. Other imaging procedures were completed during these scanning sessions and will be reported elsewhere. After scanning procedures were completed, participants were accompanied by two study team members and immediately transported back to the CPCR and remained at the CPCR until the drug effects subsided.

#### 2.2.4 Subjective-Effects Measures

Immediately following each task-free fMRI scan, participants rated the strength of subjective drug effects experienced during that scan from 0 (“none”) to 10 (“strongest imaginable”). Subjective effects ratings were verbally prompted by and then recorded by a study team member using the MRI intercom while the participant was in the scanner. Participants confirmed that they understood the meaning of each rating before the task-free fMRI scan. The overall strength of psilocybin-like effects was rated first (overall psilocybin effect). Participants then rated three measures related to mindfulness: nowness (feeling of being in present moment), letting go (degree to which person was able to let go of control of the experience), and equanimity (feeling of being in emotional balance). Participants also rated subjective effects derived from the mystical experience questionnaire (MEQ) (Barrett et al., 2015; MacLean et al., 2012): pure being and pure awareness, fusion of your personal self into larger whole, sense of reverence or sacredness, timelessness (lack of sense of space/time), ineffability (inability to explain experience with words), feelings of joy, feelings of peace and tranquility. The average score from the MEQ effects was aggregated into a single score to obtain a brief MEQ (bMEQ) score. Lastly, participants rated two dimensions of general emotional experience: positive emotional valence and negative emotional valence.

### 2.3 Preprocessing and Template ICA Modeling

#### 2.3.1 fMRI Minimal Preprocessing for Template Map Generation

The fMRI datasets were minimally preprocessed using SPM 12 (https://www.fil.ion.ucl.ac.uk/spm/) and custom code written in MATLAB (https://github.com/KKI-CNIR/CNIR-fmri_preproc_toolbox). Resting-state scans were slice time corrected using the slice in the middle of the TR as reference and realignment parameters were estimated to adjust for motion. Spatial normalization was performed by normalizing the volume in the middle of the scan to the Montreal Neurological Institute (MNI) EPI template (Calhoun et al., 2017) and linear trends were removed. Spatial smoothing was not performed so as to maximize the spatial resolution in the thalamus for template generation.

#### 2.3.2 Template ICA Model

Template ICA is a Bayesian statistical procedure that relies on empirical priors, or “templates”, which are estimates of the mean and between-subject variance of a given set of independent components for a population. We used the baseline resting-state data (N = 38 subjects) acquired during *Phase 1* of the study to generate the necessary templates to improve estimation of subject-level spatial maps derived from *Phase 2* study data. These spatial maps consist of ICA component loadings, which represent the intensity association with a particular component or network at each voxel. The template ICA model allows greater flexibility to deviate from the population mean in estimating component loading for voxels with large between-subject variance (unstable across subjects) and less flexibility for voxels with low between-subject variance (stable across subjects). The template ICA procedure is more fully explained elsewhere (Mejia et al., 2020) and is therefore only briefly discussed below.

To create the template mean and variance maps necessary to apply template ICA to *Phase II* data, we first applied spatially constrained group ICA to the thalamic region of the 38 *Phase 1* scans. First, minimally preprocessed baseline rs-fMRI data from *Phase 1* (N = 38 subjects) were masked to only include voxels inside of the thalamus using an explicit whole-thalamus mask that was extracted from the probabilistic Harvard-Oxford atlas (Desikan et al., 2006). A conservative threshold of at least 50% population-wise likelihood of intrathalamic voxel identity in the atlas was used to generate this mask, which contained 2,268 voxels. Group ICA of fMRI Toolbox (GIFT v3.0b: https://trendscenter.org/software/gift/; Medical Image Analysis Lab, Albuquerque, New Mexico) (Calhoun et al., 2001; Erhardt et al., 2011) was then used to decompose intrathalamic voxels into temporally coherent, spatially independent components. The number of thalamic independent components (TICs) to estimate for the group was selected using the maximum dimension estimate across participants (model order = 7) using an information theoretic approach. Group ICA was repeated 100 times on the baseline data with random initial conditions to assess reliability of the components (Himberg et al., 2004). Second, the timeseries of each subject’s fMRI data (N = 38 subjects) was split in half into two pseudosessions, and dual regression (DR) was used to obtain noisy estimates of subject-level TICs separately for each pseudo-session to estimate the within-subject variance. The mean and variance across all subjects and pseudo-sessions for each TIC was then calculated, and within-subject variance was removed from the total variance to estimate the between-subject variance. The “templates” or priors consist of the mean component loading and between-subject variance calculated from this step, which are output as aggregate spatial maps for each TIC. The code for template estimation and template ICA model fitting is available for Matlab (https://github.com/mandymejia/templateICA) and R (https://github.com/mandymejia/templateICAr).

The template ICA model was fit using the fMRI data from the 18 subjects that had usable task-free fMRI data from both the placebo and psilocybin sessions. The mean and variance template maps (priors) generated for each TIC in the previous step from the thirty-eight subjects’ baseline fMRI data in *Phase 1* were applied in the template ICA model (Mejia et al., 2020) to generate subject-level TICs (posterior means and standard errors) for both placebo and psilocybin sessions.

### 2.4 Effects of Psilocybin on Intrathalamic Components and Associations with Subjective Effects Measures

#### 2.4.1 Quantifying Intrathalamic Component Spatial Engagement

For each subject-and-session-specific TIC estimated via template ICA, voxelwise t-tests were performed to identify voxels showing statistically significant session-specific differences in TIC “engagement” as compared to baseline. Engagement is defined as the number of voxels with TIC estimates demonstrating a significant difference from the group mean (“template”) TIC loading at baseline, based on the posterior mean and standard error generated from template ICA. Multiplicity correction was performed by controlling the false discovery rate (FDR) at 0.01 using the Benjamini-Hochberg procedure (Benjamini and Hochberg, 1995). A conservative FDR threshold was selected to limit false positives among the large number of tests being performed (*n* = 2268 voxels within the thalamus).

After quantifying engagement for each subject and session, two approaches were used to quantify the effect of session (placebo vs. psilocybin) on TIC engagement:

1. *Engaged voxel counts*: For each TIC (TIC01-TIC07) the number of significantly engaged voxels within the whole thalamus was compared between sessions (placebo vs. psilocybin) using non-parametric Wilcoxon paired two-sided *t*-tests), adjusting for multiple comparisons (FDR = 0.05). This approach counted all significantly engaged voxels regardless of whether the voxels robustly contributed to the mean TIC loading in the baseline aggregate spatial maps. Therefore, we also counted those significant voxels that additionally showed robust contribution to the given TIC, having a z-statistic ≥ 2, and compared the number of engaged voxels between sessions using non-parametric Wilcoxon paired two-sided t-tests (see supplementary file).
2. *Probabilistic Maps of Engagement*: In addition to quantifying within-subjects effects of session type on TIC engagement, we also generated a probabilistic effect map to spatially localize those voxels showing a meaningful between-session effect at the group level. To generate the probabilistic effect maps, subject-specific TIC *engagement maps* were binarized (1 = significant engagement; 0 = no engagement), and then the binarized *engagement maps* were averaged across subjects separately for psilocybin and placebo sessions to create *mean engagement maps* for each session. Next, the difference between the psilocybin and placebo *mean engagement maps* produced a single *probabilistic effect map* that represented the proportion of subjects in which TIC engagement differed between placebo and the psilocybin sessions (*probabilistic effect map* = *placebo mean engagement map -psilocybin mean engagement map*). Voxels were then identified where at least 25% of participants showed more engagement during the placebo session as compared to the psilocybin session (no voxels showed more engagement during psilocybin compared to placebo). The 25% threshold was selected as an aggregate measure to identify those voxels showing the observed effect (psilocybin < placebo) consistently across a meaningful subset of the group (at least 5 of the 18 participants).

#### 2.4.2 Anatomic identification of between-session effects on TIC engagement

Voxel-wise overlap was calculated for the between-session probabilistic effect maps and a group of predetermined ROIs representing known subthalamic nuclei obtained from the histologically derived stereotactic Morel Atlas (Morel et al., 1997). Percent overlap represents the percentage of voxels contained within the chosen effect maps that fall into the pre-defined Morel ROIs. This same approach was taken to spatially identify voxels with robust component loading at baseline (in addition to spatially identifying the effect of psilocybin-induced changed as described above).

#### 2.4.3 Associations with Subjective-Effects

The association of between-session change in engagement (psilocybin – placebo) and between-session change in all subjective effects (see 2.2.4) was examined for each TIC. Associations of interest were identified using a standard *R*-squared effect size ≥ 0.2 (accounting for 20% of explained variance).

### 2.5 Thalamocortical Functional-Connectivity and Associations with Subjective Effects Measures

Whereas the methods detailed in the previous section were selected to examine psilocybin-induced changes in the *spatial organization* of TIC engagement among intrathalamic voxels, the following section details methods used to examine psilocybin-induced alterations in both intra and extra-thalamic *connectivity* (intra-thalamic, thalamocortical, and intra-cortical connectivity). Intra-cortical connectivity was examined for large-scale resting state networks to better understand how these networks were connected after partialing-out the effects from the thalamocortical edges. A summary of procedures is outlined below:

1. *DR to obtain subject and session-specific TIC timecourses*: DR using group ICA-derived template spatial maps was applied to minimally preprocessed subject- and session-specific resting-state data to obtain subject-level TIC timecourses (TIC01-TIC07).
2. *DR to obtain subject and session-specific cortical ICA-derived network timecourses [CINs]*: DR was applied to minimally preprocessed subject- and session-specific cortical resting-state data using aggregate cortical ICA-derived networks (CIN) that were previously identified by Smith et al (Smith et al. 2009). The Smith networks include aggregate maps corresponding to three visual areas [medial (IC08); occipital pole (IC09); lateral visual (IC10)], default mode network (DMN; IC11), cerebellar network (IC12), sensorimotor network (SMN; IC13), auditory network (IC14), executive control network (IC15), and two lateralized frontoparietal networks [right FPN (IC16) and left FPN (IC17)]. Prior to DR, voxels in the probabilistic whole-thalamus mask were excluded from each network spatial map.
3. *Regress nuisance timecourses and remove outlier timepoints*: Nuisance regression was used to remove the influence of non-neural signal estimated from white matter (WM) and cerebrospinal fluid (CSF) (Behzadi et al., 2007; Muschelli et al., 2014) from TIC and CIN timecourse data. The regressors included the first two principal components from the WM and CSF as well as four discrete cosine transform bases. The resulting timecourses were despiked to remove the influence of outlier timepoints (Analysis of Functional Neuroimages: http://afni.nimh.nih.gov/afni; NIMH Scientific and Statistical Computing Core, Bethesda, Maryland).
4. *Thalamocortical connectivity matrix [7×10]*: Subject- and session-specific 7×10 partial correlation matrices were generated using ridge regression (rho = 1) (Lombardo et al., 2019), converted from *r* to *z*-statistics using Fisher’s transformation, and compared between sessions using paired two-sided t-tests to examine between-session differences in both intra-thalamic, thalamocortical, and intra-cortical connectivity (FDR = 0.05).
5. *Associations with Subjective-Effects*: Between-session changes in functional connectivity were associated with between-session changes in subjective-effects (see *section 2.4.3*) using Spearman rank correlations. Comparisons were limited to those thalamocortical and intra-cortical connectivity pairs that showed significant between-session differences in functional connectivity. Associations explaining 20% of observed variance between sessions were reported (R^2^ ≥ 0.2).

### 2.6 Whole-Thalamus Seed Connectivity with Cortical ICA-derived Networks

To examine the impact of our thalamic parcellation on our results and for comparison to the existing literature, we calculated thalamocortical connectivity using the mean timecourse across all thalamic voxels. Each voxel timecourse was nuisance regressed using the approach described in section 2.5 and then averaged to obtain a single seed timecourse of the whole thalamus. The whole-thalamus seed timecourse was then correlated with the cortical ICA-derived networks (CINs) that were obtained using DR as described in section 2.5. These correlations were Fisher *z*-transformed and paired two-sided t-tests were performed to examine between-session (placebo-psilocybin) differences in whole-thalamus thalamocortical connectivity (FDR = 0.05).

## 3. RESULTS

### 3.1 Identification of Spatially and Functionally Independent Intrathalamic Components

#### 3.1.1 Visualization of Intrathalamic Components at Baseline Shows Spatial Clustering and Bilateral Symmetry

Two approaches were taken to examine the loading of intra-thalamic voxels onto each TIC. First, we used a winner-take-all assignment strategy using mean component loadings at baseline for each voxel, which resulted in spatially clustered regions demonstrating a reasonable degree of bilateral symmetry (lighter shading, Figure 1). A separate but complementary approach was then taken to identify voxel-to-component assignments. All thalamic voxels were thresholded for each TIC to select voxels demonstrating robust mean component loading at baseline (z ≥ 2; darker shading, Figure 1). In this case, there were no instances of any individual voxel demonstrating robust loading in multiple TICs, suggesting complete spatial segregation of robustly contributing voxels. These voxels also demonstrated spatial clustering and reasonable bilateral symmetry, providing evidence for construct validity; further, these voxels were contained within the larger group of voxels that were assigned to each TIC utilizing the more inclusive winner-take-all assignment strategy (lighter shading, Figure 1).

**Figure 1:**
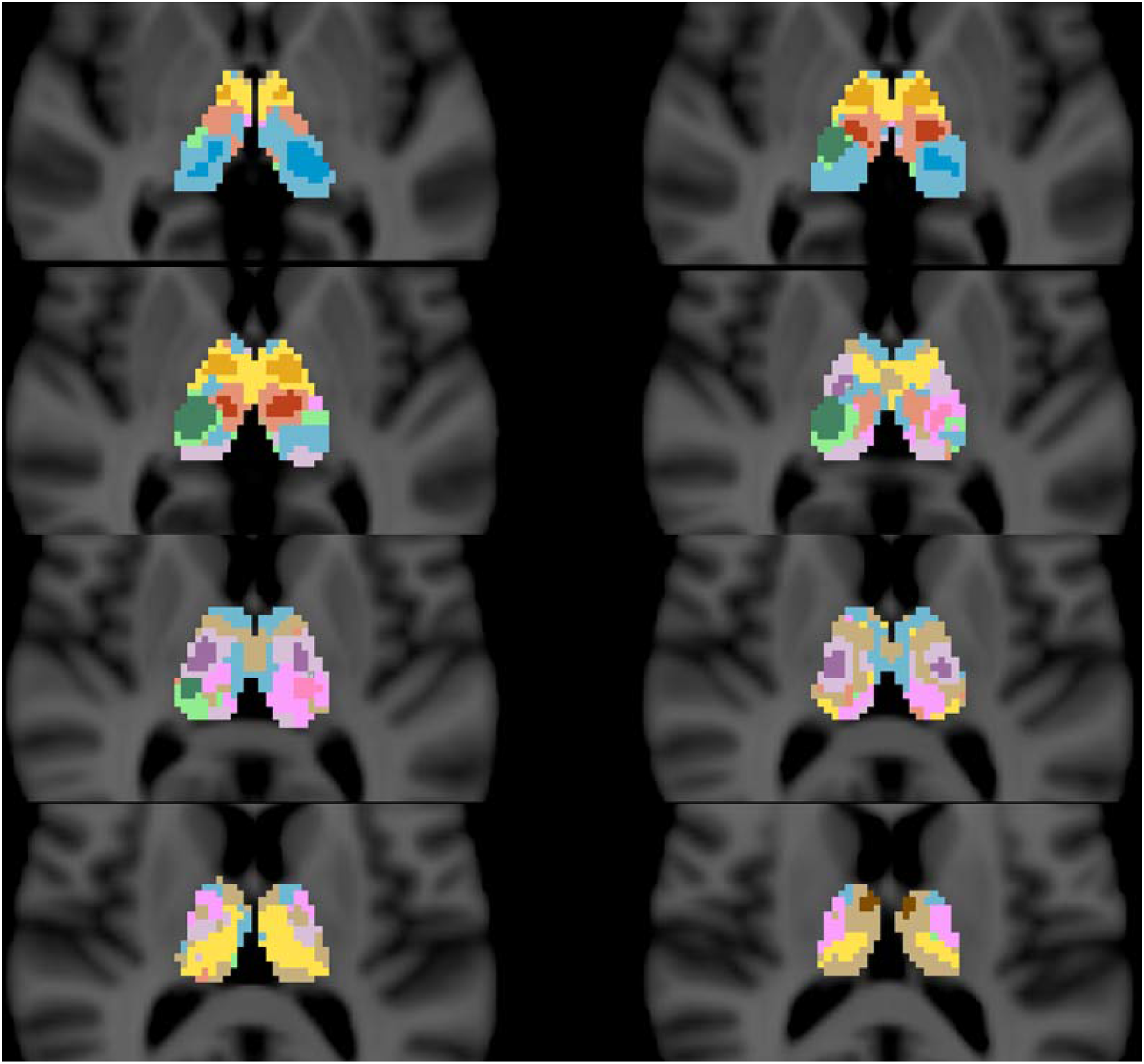
Visual display of thalamic parcellation derived from spatially constrained group Independent Component Analysis (ICA) applied to the Phase I data, comparing two voxel-wise assignment strategies. Each panel plots Thalamic Independent Component (TIC) assignment for all visible intra-thalamic voxels on an axial slice of the CH-best template from MRIcron. Lighter shades represent a more inclusive winner-take-all assignment strategy, in which every thalamic voxel is assigned to the component with the highest loading at baseline. Darker shades represent only voxels with robust component loading at baseline (z ≥ 2). Components are labeled in two shades (one for each assignment strategy) of each of the following colors: TIC01- brown, TIC02- green, TIC03- blue, TIC04- orange, TIC05- red, TIC06- pink, TIC07- purple. The visualization shows that components have larger and more weakly contributing peripheral areas that encompass smaller cores with robust contribution. For each component, both areas are clearly spatially contiguous and bilaterally symmetrical.

#### 3.1.2 Intrathalamic Components at Baseline Demonstrate Distinct Patterns of Overlap with Intrathalamic Nuclei

The results above align with our understanding of the thalamus as a structure comprised of bilaterally symmetrical and functionally distinct nuclei. In order to further illustrate this, we calculated the percent overlap of voxels with robust mean loading at baseline for each TIC (z ≥ 2; darker shading, Figure 1) with known histologically-derived ROIs from the Morel Atlas (Morel et al., 1997). Each TIC demonstrates a unique pattern of spatial overlap with intrathalamic nuclei, with most demonstrating robust (>50%) spatial overlap with at least one ROI, and each TIC demonstrating best fit to a unique Morel ROI (TIC1-LD/LP, TIC2-VP, TIC3-PumM, TIC4-VA, TIC5-MD, TIC6-PuA, TIC7-VL; Figure 2).

**Figure 2:**
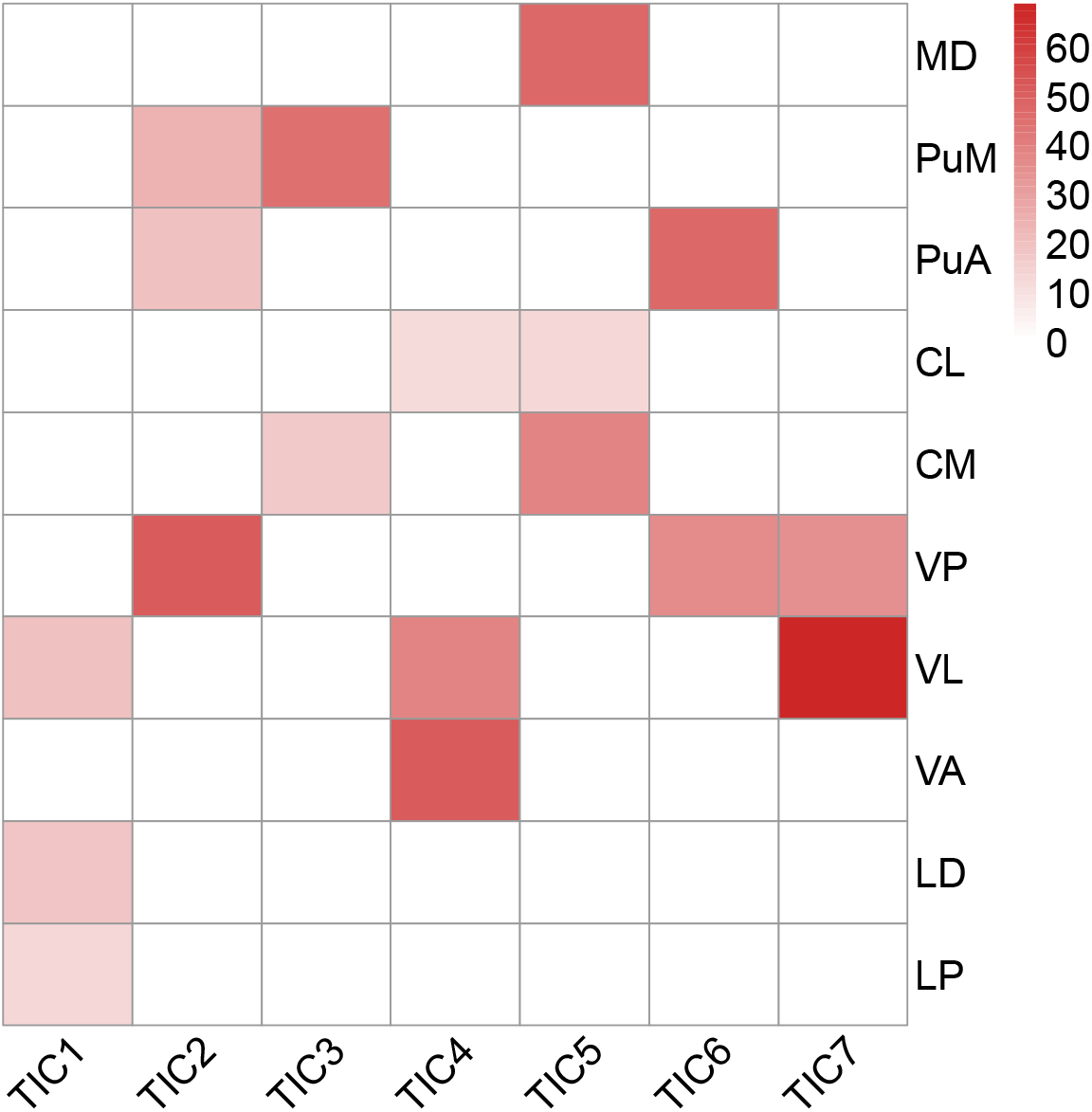
Percent overlap between histologically identified thalamic nuclei (Morel et al., 1997) and intrathalamic components thresholded at z ≥ 2. Only those nuclei with ≥10% overlap for any TIC are displayed. MD = mediodorsal; PuM = medial pulvinar; PuA = anterior pulvinar; CL = central lateral; CM = central medial; VP = ventral posterior; VL = ventral lateral; VA = ventral anterior; LD = lateral dorsal; LP = lateral posterior.

### 3.2 Altered spatial organization of thalamic independent components during psilocybin vs. placebo administration

#### 3.2.1 Thalamic Independent Components Show Decreased Engagement during Psilocybin vs. Placebo Administration

Overall engagement was significantly lower during psilocybin (PSI) sessions than during placebo (PLA) sessions for TIC03 (*p* = 0.006; PLA > PSI) and TIC05 (*p* = 0.01; PLA > PSI) (FDR = 0.05) (Figure 3). Considering only voxels in each TIC demonstrating robust mean component loading at baseline (z ≥ 2), TIC02 (*p* = 0.03; PLA > PSI) and TIC07 (*p* = 0.02; PLA > PSI) also showed significantly decreased engagement during psilocybin sessions, in addition to TIC03 (*p* = 0.02; PLA > PSI) and TIC05 (*p* = 0.03; PLA > PSI), (FDR = 0.05) (Figure S1).

**Figure 3:**
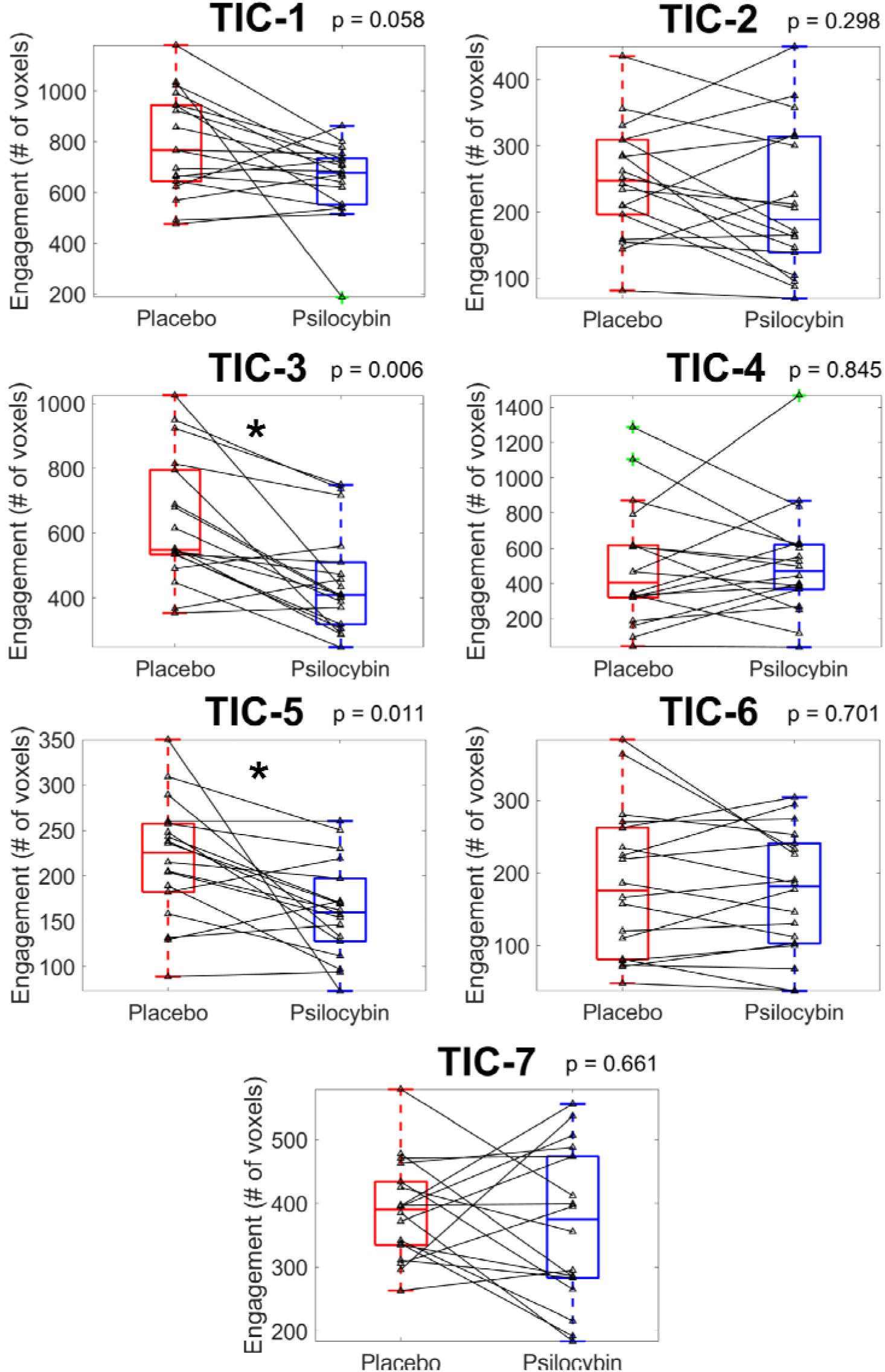
Overall engagement (number of voxels) for each Thalamic Independent Component (TIC01 to TIC07) in each drug condition. Non-parametric Wilcoxon two-sided paired t-test p-values for the comparison of voxel counts between placebo and psilocybin sessions are displayed at the top right-hand corner of each panel, corrected at FDR = 0.05. Engagement within only voxels demonstrating robust component loading at baseline (z ≥ 2) was also significantly decreased for TIC03 and TIC05, consistent with the whole thalamus voxel count results displayed in Figure 2, with additional components TIC02 and TIC07 also showing significantly decreased engagement during the psilocybin vs. placebo session. These results are displayed in the supplementary material (Figure S1).

#### 3.2.2: Effect of Psilocybin on Intrathalamic Functional Organization Localized to Medial Pulvinar and Mediodorsal Nuclei (psilocybin < placebo)

The thresholded between-session effect maps from the four TICs that showed significant between-session (PSI vs. PLA) differences in intrathalamic engagement (TICs 02, 03, 05, 07) were compared to histologically derived ROIs of thalamic nuclei (Morel et al., 1997). Differences in engagement in each of these four TICs were primarily localized to either the Mediodorsal or Medial Pulvinar nuclei (Figure 4).

**Figure 4:**
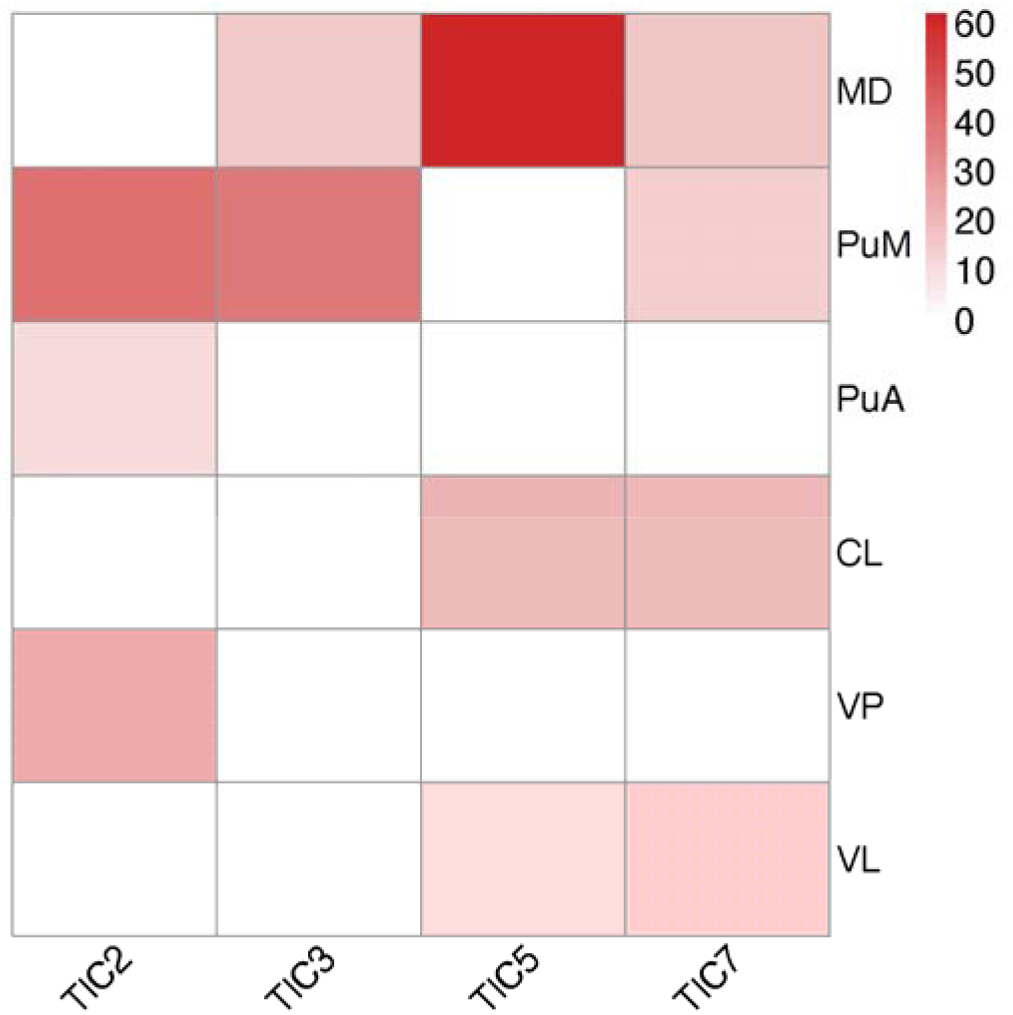
***Left*** - Percent overlap between histologically identified Morel thalamic nuclei and intrathalamic component *probabilistic effect maps* showing significant decreases in engagement during psilocybin vs. placebo. Only those nuclei with ≥10% overlap are displayed. MD = mediodorsal; PuM = medial pulvinar; PuA = anterior pulvinar; CL = central lateral; VP = ventral posterior; VL = ventral lateral.

#### 3.2.3 Between-Session Differences in Thalamic Engagement and Associations with Subjective Effects

A total of six associations between the change in engagement voxel counts (placebo-psilocybin) and subjective-effects (placebo-psilocybin) met the established criterion for a very-large association (Funder and Ozer, 2019) (*r*^2^ ≥ 0.2 or *r* > 0.45) using Spearman rank correlation. Participants showing larger decreases in engagement of TIC03 during psilocybin sessions compared to placebo had larger subjective effect ratings of “letting go” (*r* = −0.47). Participants showing increased engagement of TIC07 during psilocybin sessions had larger subjective effects of “equanimity” (*r* = 0.55), “pure being” (*r* = 0.52), “joy” (*r* = 0.52), and an aggregate measure of “mystical experience” (bMEQ) (*r* = 0.54) during psilocybin sessions. Further, participants showing larger decreases in TIC06 engagement during the psilocybin vs. placebo session showed larger increases in feeling of “joy” during the psilocybin session (*r* = −0.48) (Figure 5).

**Figure 5:**
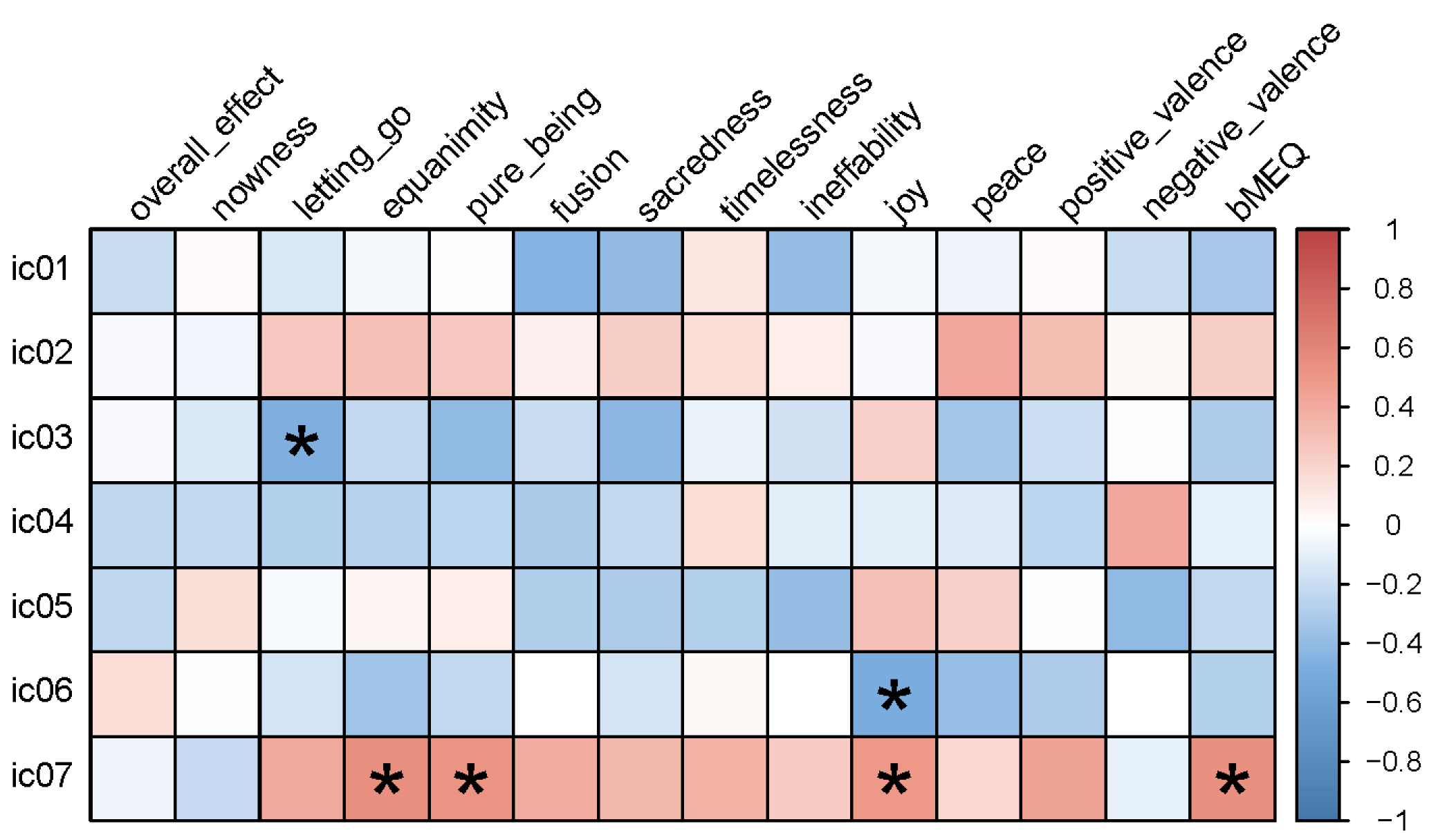
Spearman rank correlations between change in engagement voxel counts and change in subjective effects from placebo to psilocybin. Correlations with *r*^2^≥0.20 (meeting criteria for a very large association) are indicated with an asterisk*.

### 3.3 Altered between-network connectivity distributed across thalamus-derived ICA and cortical-derived ICA networks

#### 3.3.1 Predominant Decreases in Functional Connectivity During Psilocybin vs. Placebo Sessions

Overall, during psilocybin sessions thirteen edges showed significantly decreased connectivity compared to only five edges showing significantly increased connectivity. For *intrathalamic connectivity*, two edges showed significantly decreased connectivity and one edge showed significantly increased connectivity during psilocybin sessions. For *thalamocortical connectivity*, five edges showed significantly decreased connectivity whereas one edge showed significantly increased connectivity during psilocybin sessions. For *intra-cortical connectivity*, six edges showed significantly decreased connectivity and three edges showed significantly increased connectivity during psilocybin sessions (Figure 6; Table 1).

**Figure 6:**
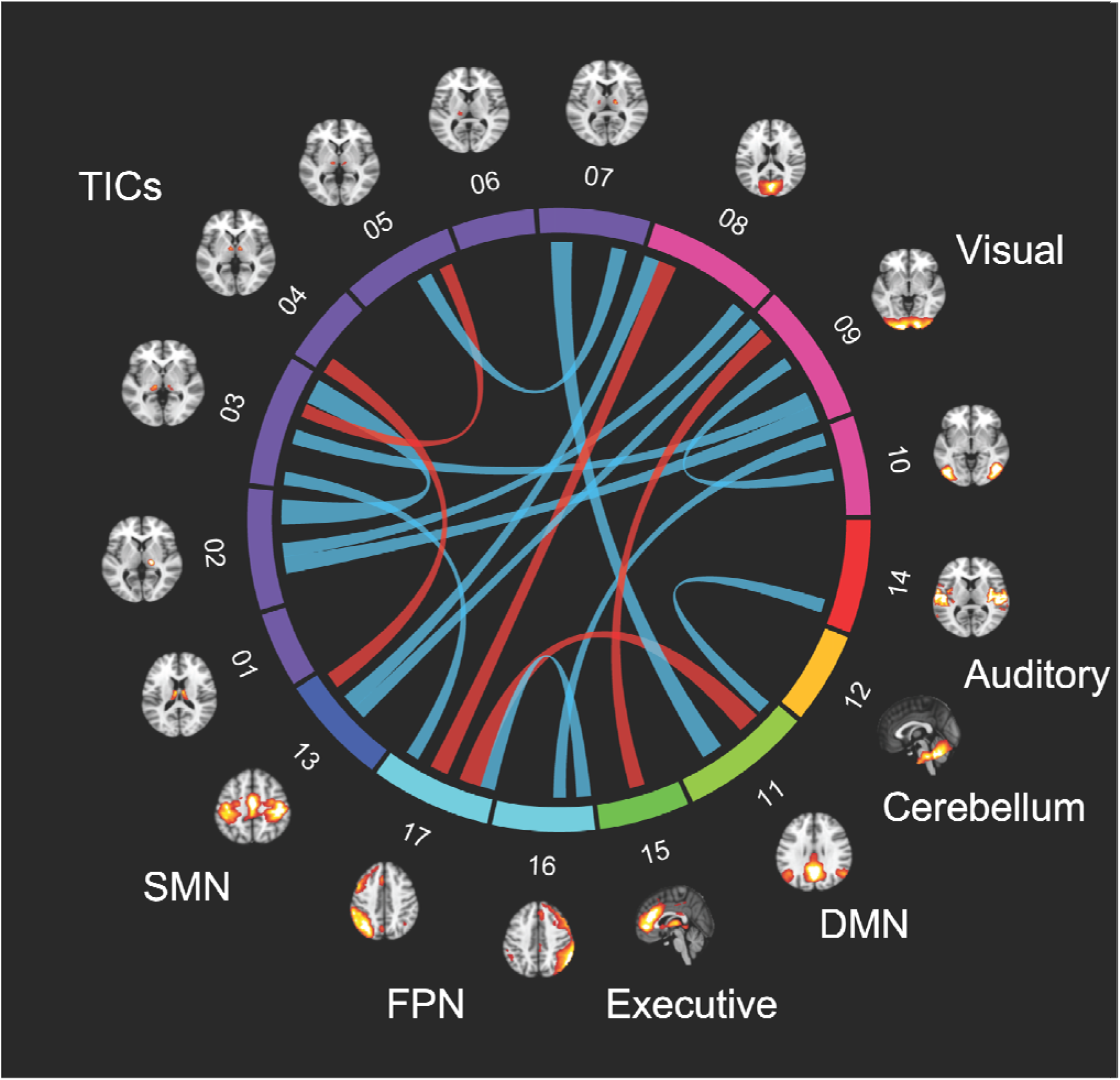
Changes in between-component (and between-network) connectivity between placebo (PLA) and psilocybin (PSI) sessions (FDR = 0.05). Red lines between networks indicate increased connectivity on psilocybin (PLA < PSI) and light blue lines indicate decreased connectivity on psilocybin (PLA > PSI). Axial or sagittal slices outside of the circumference of the connectogram depict components included in the connectogram, and color codes on the circumference of the connectogram denote the overall network sets to which each component belongs. Purple: Thalamic Independent Components (TIC01 to TIC07). Pink: visual network (IC-08, IC-09, and IC-10). Red: auditory network (IC-14). Yellow: cerebellar network (IC-12). Chartreuse: Default Mode Network (“DMN”, IC-11). Spring Green: Executive Control Network (“Executive”, IC-15). Cyan: Right/Left Frontoparietal Networks (“FPN”, IC-16 and IC-17). Dark Blue: Somatosensory network (“SMN”, IC-13). Results using a correction level of FDR = 0.01 are shown in the Supplement (Figure S2).

**Table 1:** Edges showing significant between-session differences (FDR = 0.05). The top panel shows intra-thalamic connectivity (TIC01-TIC07) and thalamocortical connectivity (TIC01-IC17). The bottom panel shows significant edges between ICA-derived cortical networks-only (IC08-IC17). Positive t-statistics represent decreased connectivity on psilocybin (PLA > PSI) and are displayed in black; negative t-statistics represent increased connectivity on psilocybin (PLA < PSI) and are displayed in red.

|  | Significant Edges (FDR = 0.05) |  | T statistics<br>[placebo-psilocybin] | FDR corrected *p*-values |
| --- | --- | --- | --- | --- |
| THALAMIC CONNECTIVITY | TIC-02 | TIC-03 | $t = 5.64$ | $p < 0.001$ |
| | | IC-08 | $t = 3.17$ | $p = 0.017$ |
| | | IC-09 | $t = 3.88$ | $p = 0.002$ |
| | TIC-03 | TIC-05 | $t = -2.99$ | $p = 0.023$ |
| | | IC-09 | $t = 3.34$ | $p = 0.011$ |
| | | IC-17 | $t = 2.99$ | $p = 0.023$ |
| | TIC-04 | IC-13 | $t = -3.73$ | $p = 0.004$ |
| | TIC-05 | TIC-07 | $t = 3.47$ | $p = 0.008$ |
| | TIC-07 | IC-11 | $t = 4.79$ | $p < 0.001$ |
| INTRA-CORTICAL CONNECTIVITY | IC-08 | IC-13 | $t = 3.45$ | $p = 0.008$ |
| | | IC-17 | $t = -4.13$ | $p = 0.001$ |
| | IC-09 | IC-10 | $t = 2.73$ | $p = 0.046$ |
| | | IC-13 | $t = 3.00$ | $p = 0.023$ |
| | | IC-15 | $t = -3.48$ | $p = 0.008$ |
| | IC-10 | IC-16 | $t = 2.89$ | $p = 0.03$ |
| | IC-11 | IC-14 | $t = 3.12$ | $p = 0.018$ |
| | | IC-17 | $t = -4.94$ | $p < 0.001$ |
| | IC-16 | IC-17 | $t = 3.28$ | $p = 0.012$ |

#### 3.3.2 Between-Network Connectivity and Associations with Subjective Effects

Changes in between-network connectivity (BNC) from placebo to psilocybin was examined between each TIC (TIC01 to TIC07) and each cortical ICA-derived network (IC08-17) (Figure 7).

1. *Intra-thalamic BNC*: Participants showing larger decreases in connectivity between TIC02-TIC03 during psilocybin sessions showed larger decreases in “fusion” (*r* = 0.47), “sacredness” (*r* = 0.53), and “peace” (*r* = 0.49). Further, participants showing larger decreases in connectivity between TIC05-TIC07 during psilocybin sessions showed larger decreases in “timelessness” (*r* = 0.54) and “joy” (*r* = 0.48) (Figure S3).
2. *Thalamocortical BNC*: Participants showing larger decreases in connectivity between TIC02-IC08 during psilocybin sessions showed larger increases in “positive-valence” (*r* = −0.57). Participants showing larger decreases in connectivity between TIC03-IC09 during psilocybin sessions showed larger increases in “overall effect” (*r* = −0.47). TIC03-IC17 showed larger decreases in connectivity during psilocybin being associated with larger decreases in “negative valence” (*r* = 0.45) and increases in “joy” (*r* = −0.55). Finally, participants showing larger decreases in connectivity between TIC07-IC11 during psilocybin showed larger decreases in “letting go” (*r* = 0.50) and larger increases in “negative valence” (*r* = −0.50) (Figure S4).
3. *Intra-cortical BNC*: Participants showing larger *increases* in connectivity between IC08-IC17 during psilocybin sessions showed larger decreases in “timelessness” (*r* = −0.61). Participants showing larger decreases in connectivity between IC11-IC14 during psilocybin sessions showed larger increases in “nowness” (*r* = −0.46). Finally, participants showing larger decreases in connectivity between IC16-IC17 during psilocybin sessions showed larger increases in “overall effect” (*r* = −0.45), “fusion” (*r* = −0.47), “sacredness” (*r* = −0.46), and “ineffability” (*r* = −0.53) (Figure S5).

**Figure 7:**
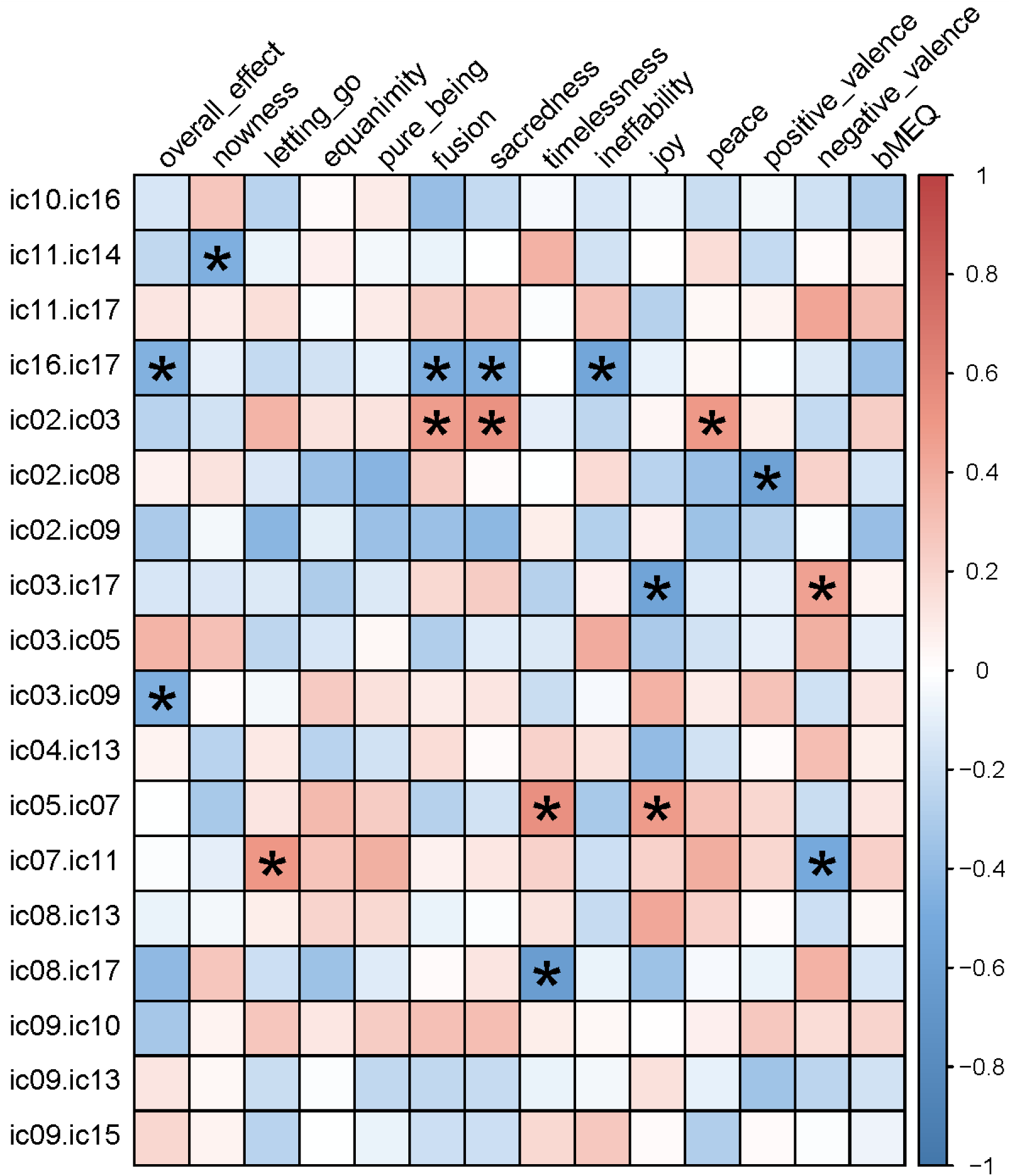
Changes in the strength of significant edges (FDR = 0.05) are associated with changes in subjective experience from placebo to psilocybin (shown using Spearman rank correlations). Correlations with *r*^2^ ≥ 0.20 are highlighted with an asterisk*. IC01-07: Thalamic Independent Components (“TIC”s), IC08-10: Visual Network, IC11: Default Mode Network (“DMN”), IC12: Cerebellar Network, IC13: Somatosensory Network (“SMN”), IC14: Auditory Network, IC15: Executive Control Network, IC16-17: Right/Left Frontoparietal Networks (“FPN”).

#### 3.3.3 No Significant Thalamocortical Connectivity Edges Using Whole-Thalamus Seed Approach

No between-session differences in whole-thalamus seed connectivity with the cortical ICA-derived networks were observed after correcting for multiple comparisons (FDR = 0.05). The largest change in connectivity between the whole thalamus seed and cortical networks was observed with the occipital-pole (IC09), showing a marginally greater connectivity during psilocybin sessions (*t* = −2.68; FDR corrected *p* = 0.07). Further, other ICs that showed large but non-significant increases in functional connectivity with the whole thalamus seed during psilocybin included the DMN (IC11; *t* = −2.23; FDR corrected *p* = 0.08) and left FPN (IC17; *t* = −2.11; FDR corrected *p* = 0.08), with the SMN (IC13; *t* = 2.3; FDR corrected *p* = 0.08) showing more negative connectivity with the whole thalamus seed.

The direction of between-session connectivity differences using the seed-based approach were opposite those observed for the significant TIC thalamocortical edges. One possible explanation for the differences between the seed-based and ICA approaches are that psilocybin effects might be localized to small thalamic regions that are washed out when all voxels are considered equally. To visualize this effect, we calculated correlations between all thalamic voxel timecourses and both DMN (IC11) and occipital-pole (IC-09) cortical ICs. Both the DMN and occipital-pole showed increased connectivity during the psilocybin session using the seed-based approach but showed decreased connectivity during the psilocybin session using the ICA-based approach (i.e., DMN to TIC07; occipital-pole to TIC02; occipital-pole to TIC03). When examining between-session differences in thalamocortical voxelwise connectivity with the DMN timecourse, a region localized to TIC07 showed the effect of decreased connectivity on-psilocybin, whereas voxels outside this region showed the opposite effect. The same observation was made for the occipital-pole, where regions localized to both TIC02 and TIC03 showed the effect of decreased connectivity on-psilocybin, whereas voxels outside these ROIs showed the opposite effect (Figure 8). The observations displayed in Figure 8 provide an explanation for why the seed-based approach produced opposite findings as compared to the ICA-based approach. Collectively, these findings represent an advantage of ICA over seed-based approaches by providing a more nuanced evaluation of thalamocortical connectivity changes that occur during psilocybin administration.

**Figure 8:**
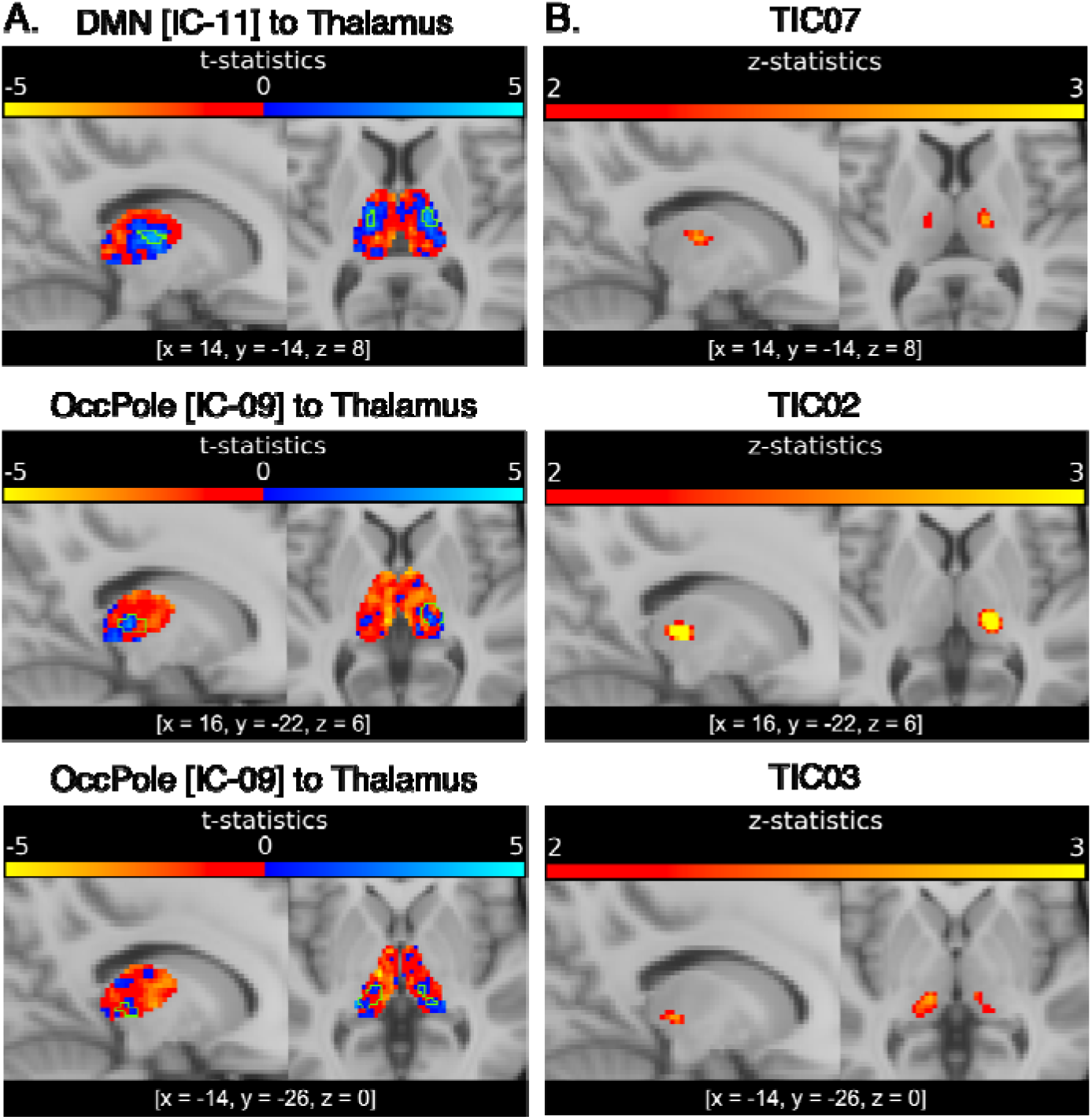
Voxelwise connectivity between all thalamic voxels (n = 2268) and both the Default Mode Network (“DMN”, IC11) and Occipital Pole (IC09). (Panel A) Voxelwise thalamocortical connectivity t-statistics for DMN and Occipital Pole cortical ICs. Blue = placebo > psilocybin. Red = placebo < psilocybin. Green outline shows area of voxels located within TICs shown on corresponding row in panel B. (Panel B) Mean TIC engagement from baseline data thresholded at z ≥ 2. Shown using neurological convention.

## 4. DISCUSSION

Using a novel data-sparing ICA-based approach, we report several significant psilocybin-induced changes in both intra- and extra-thalamic patterns of functional organization and connectivity not reported in previous studies that treat the thalamus as a unitary structure. Given the known functional and structural segregation of thalamic nuclei, ICA has been increasingly employed in order to generate functional subdivisions of the thalamus. Congruent with several existing studies (Tian et al., 2020; Zhang and Li, 2017), our results support the utility of ICA in generating parcellations of the thalamus from baseline resting state data that show functional distinction between components and also clear spatial clustering, especially for voxels with robust contribution to each component. Using this approach, we compared thalamic functional parcellations generated before vs after the acute administration of psilocybin, which to our knowledge, is the first such report.

The acute administration of psilocybin caused significant alterations in the spatial organization of functionally derived intrathalamic components, with several components showing a significant decrease in the number of engaged voxels after administration of psilocybin. Significant changes in thalamic connectivity with cortical networks were also observed, showing predominantly decreases in connectivity between intrathalamic components and cortical networks. Given that both cortical (Smith et al., 2009) and subcortical (Tian et al., 2020) networks are highly stable in the resting state, and given that psilocybin is known to cause acute alterations in global functional connectivity (Preller et al., 2020), our results suggest that psilocybin may substantially alter both intrathalamic and thalamocortical connectivity in a spatially constrained fashion, with the principal effect of focally reducing intrathalamic and thalamo-cortical connectivity. Our results also show that several intrathalamic and thalamocortical alterations in connectivity are also associated with reported acute subjective effects, which is especially significant given that certain reported subjective effects have been shown to predict the therapeutic efficacy of psilocybin (Griffiths et al., 2018)

Through the use of a novel data-sparing approach, our study reveals distinct focal changes in intrathalamic functional organization that are associated with the acute administration of psilocybin, are apparent at 3T resolution, are associated with both alterations in thalamocortical connectivity and subjective drug effect, and are spatially aligned with nuclei previously implicated in the psychedelic state. Thalamic voxels most impacted by the administration of psilocybin were spatially clustered, and when compared to a histologically derived atlas, were largely localized to the mediodorsal and pulvinar nuclei. Localization primarily to these two nuclei is consistent with evidence that, within the thalamus, the mediodorsal and pulvinar nuclei maximally express the 5-HT_2AR_ receptor (Wai et al., 2011). The findings are also consistent with those from animal models revealing the mediodorsal nucleus to be implicated in the acute effects of LSD (Inserra et al., 2021). While ours are the first reported findings regarding the effects of psilocybin on the functional organization of intrathalamic nuclei in humans, findings from one fMRI study after LSD administration did reveal distinct spatial changes in thalamocortical connectivity (Müller et al., 2018).

In addition to observed psilocybin-associated alterations in the internal functional organization of the thalamus, patterns of extrathalamic connectivity were also significantly altered after administration of psilocybin, with specific components showing decreased connectivity with large-scale cortical networks. Only one previous study reported on thalamocortical connectivity after acute administration of psilocybin; treating the thalamus as a single functional unit, this study reported increased connectivity between the thalamus and a “task positive” network (Carhart-Harris et al., 2013). Interestingly, our findings of focal component-specific decreases in connectivity are in seeming contrast with those from this 2013 study and several other fMRI studies examining the effect of LSD that also use whole-thalamus approaches and report apparent increases in thalamocortical connectivity (Müller et al., 2017; Tagliazucchi et al., 2016). If we employ a similar whole-thalamus approach using this current dataset, we also find overall numerical increases in thalamo-cortical connectivity, with the largest increases being observed with the visual network, although these changes notably did not reach statistical significance.

Our study is the first to report on the engagement and connectivity of specific thalamic subregions during acute administration of psilocybin. While our findings of decreased thalamo-cortical connectivity, particularly with visual and default mode networks, seemingly contrast with previous studies that report primarily increased connectivity using a whole-thalamus approach (Müller et al., 2017; Tagliazucchi et al., 2016), they are consistent with several more recent reports on LSD-induced changes in thalamocortical connectivity, which suggest nuanced regional effects including both increases and decreases in functional connectivity (Preller et al., 2020, 2019, 2018). It therefore seems possible that whole-thalamic masking may be less effective than ICA-based parcellation at detecting certain focal patterns of altered thalamic engagement and connectivity that occur during acute exposure to psilocybin and perhaps other classic psychedelics, likely reflective of the functional differences between specific intrathalamic nuclei.

In summary, our results demonstrate that using an average, whole-thalamus seed, psilocybin appears to be associated with modest increases in thalamo-cortical connectivity, which is consistent with prior whole-thalamus studies of both psilocybin and LSD effects. In contrast, our novel interrogation of functionally derived intrathalamic components reveals focal *decreases* in connectivity that are spatially clustered. Observed changes in both intra- and extra-thalamic connectivity are largely specific to thalamic nuclei (mediodorsal and pulvinar nuclei) and cortical networks (visual and default mode) that express 5-HT_2AR_ receptors and are implicated in the acute effects of the classic psychedelics. Further, in several instances, altered thalamocortical connectivity was associated with reported acute subjective effects with potential clinical relevance.

Coarse measures of thalamocortical connectivity, obtained using a whole-thalamus seed, may obscure powerful nuanced nuclei-specific effects associated with the acute administration of psilocybin. While whole-thalamus masking is likely often employed due to limitations in sample size and resolution, our results demonstrate that the use of a data-sparing approach to create functional thalamic subdivisions can reveal focal underlying effects during acute drug exposure. The potential application of this methodology to higher-resolution data, other imaging modalities, or during acute exposure to other psychoactive substances and non-pharmaceutical altered perceptual states would all be useful future approaches in fully characterizing and understanding the significance of these reported findings.

## ACKNOWLEDGEMENTS

This work was supported by a grant from the Heffter Research Institute (R.R.G.) and by the financial support of the Johns Hopkins Center for Psychedelic and Consciousness Research which was funded by the Steven and Alexandra Cohen Foundation, Tim Ferriss, Matt Mullenweg, Blake Mycoskie, and Craig Nerenberg.

## Notes

https://github.com/mandymejia/templateICA

https://github.com/mandymejia/templateICAr

https://github.com/KKI-CNIR/CNIR-fmri_preproc_toolbox

### Competing Interest Statement

RRG is a board member of the Heffter Research Institute. No other authors have competing interests.

